# Cooperative foraging behavior during larval stage affects fitness in adult *Drosophila*

**DOI:** 10.1101/2020.05.04.076869

**Authors:** Mark Dombrovski, Rives Kuhar, Alexandra Mitchell, Hunter Shelton, Barry Condron

## Abstract

Cooperative behavior can confer advantages to animals. This is especially true for cooperative foraging which is thought to provide fitness benefits through more efficient acquisition and consumption of food. While examples of group foraging have been widely described, principles governing the formation of such aggregations as well as rules that determine cooperative group membership remain poorly understood. Here we take advantage of an experimental model system featuring cooperative foraging behavior in *Drosophila*. Under crowded conditions, fruit fly larvae form coordinated digging groups (clusters), in which individuals are linked together by sensory cues and stable group membership requires prior social experience. However, fitness benefits of *Drosophila* larval clustering remain to be determined. We demonstrate that when grown in crowded conditions on food that has been partially processed by other larvae, animals experience a developmental delay presumably due to the lesser nutrient value of the substrate. Intriguingly, these same conditions promote the formation of cooperative foraging clusters which further extends the larval stage compared to non-clustering animals. Remarkably, this developmental retardation also results in a relative increase in wing size, which is a good indicator of adult fitness. Thus, we find that the clustering-induced developmental delay is accompanied by adult fitness benefits. This suggests that foraging group membership provides advantages over solitary feeding in processed food. Therefore, cooperative behavior, while delaying development, may have evolved to give *Drosophila* larvae benefits when presented with competition for limited food resources.

## Introduction

Group foraging is a major component of cooperative animal behavior (Allee 1927). It can be defined as inter- and intraspecific cooperation in search, acquisition, defense and consumption of common food resources and can provide benefits in survival and reproduction for a variety of animals (Sumpter 2005; Giraldeau and Caraco 2018). Participation in a cooperative feeding group can provide a significant enhancement in average feeding efficiency for two reasons: (1) increased food processing efficiency resulting in less investment for higher nutritional return (Valone 1989; Cash et al. 1993; Pöysä 1994; Tripet et al. 2009) and (2) a potential to sequester a common food resource from competing species or different populations of the same species (Foster 1985; Dubois 2003; Tania 2012). In addition, aggregation can also lead to a decreased risk of predation (Turchin and Kareiva 1989; Rohlfs and Hoffmeister 2004). All of these factors increase individual’s chances of survival and reproductive success which serve as the main measures of fitness (Clark and Mangel 1986). Importantly, benefits of cooperative foraging often take effect under certain conditions when availability and distribution of food resources determines advantage of cooperation, (Monaghan and Metcalfe 1985; Scheel and Packer 1991; Eklöv 1992; Amor et al. 2010). This may serve an example of Allee effect (Courchamp et al. 1999) in context of cooperative feeding, where individual’s fitness gains correlate with group size and density only to a certain limit, beyond which acquired benefits get leveled and negatively outweighed by emerging complex non-trophic factors of group membership, such as intra-group competition (Clark 1987; Cash et al. 1993; Rivers et al. 2011).

Cooperative foraging behavior has been observed among a broad range of animal taxa. Group hunting strategies were described in carnivorous mammals (Clark 1987), birds (Hector 1986) and fish (Eklöv 1992), all of which predominantly utilize active coordinated search tactics when hunting prey. Herbivores widely engage in cooperative foraging as well (Monaghan and Metcalfe 1985; Foster 1985; Pöysä 1994). Invertebrates and in particular arthropods, also display a variety of cooperative foraging behaviors including examples of interspecific aggregations that facilitate localization and consumption of food is observed among (Boulay et al. 2019, Cocroft 2005, Amor et al. 2010, Tripet et al. 2009). Many studies focus on insect larvae in which food consumption is a top priority (Fitzgerald and Peterson 1988), and animals are therefore very sensitive to trophic advantages of foraging group membership. Indeed, highly efficient foraging groups were documented in sawfly larvae (Ghent 1960; Lindstedt et al. 2019) various species of caterpillars (Clark and Faeth 1997) and even corpse-devouring necrophagous flies (Rivers et al. 2011; Scanvion et al. 2018). Importantly, factors that provide trophic benefits can serve as tradeoffs in case of severe overcrowding (e.g. overly elevated temperature and proteotoxic stress caused by excessive tissue digestion), implying a complex non-linear pattern of relationship between group size, food availability and individuals’ investment into cooperative efforts (Valone 1989; Courchamp 1999; Giraldeau and Caraco 2018). In this regard, using a laboratory model system might provide the right tools and metrics to begin dissecting out various complex parameters governing collective foraging behavior.

To address these questions, we make use of a novel experimental model system featuring cooperative foraging behavior in larval *Drosophila melanogaster*. Interestingly, although behavioral and developmental aspects of larval solitary foraging behavior were addressed long time ago (Sokolowski 1982; Godoy-Herrera 1977, 1986; Wu et al. 2003; Kim et al. 2017), mechanistic and neuroethological features of cooperative foraging in larval *Drosophila* have only recently been characterized (Durisko et al., 2014; Dombrovski et al. 2017, 2019, Khodaei and Long, 2019). Feeding clusters form in semi-liquid food, comprise of 10-200 animals and share a unique set of characteristics that make it an attractive model for studying collective social behavior. In particular, clustering larvae engage in synchronous reciprocating digging, where each group member utilizes visual cues to coordinate movements with immediate neighbors (Dombrovski et al. 2017). Intriguingly, cluster membership and ability to efficiently engage in visually guided cooperation requires prior visual and social experience during a critical period in development (Slepian et al. 2015; Dombrovski et al. 2017). In addition, emergence of clustering is associated with functional changes of larval visual circuit (Dombrovski et al. 2019). This raises the question as to the function of this behavior and its emergence in the evolution. Our study was aimed at identifying specific ecological benefits associated with social clustering, whether it refers to more efficient burrowing (Dombrovski et al. 2017), escape from predators (Carton et al. 1985) or more efficient utilization of food resources (Gregg et al. 1990; Scanvion et al. 2018).

## Results

### Processed food delays development and reduces size in *Drosophila*

Processed food and crowded conditions increase the duration of larval development (Dombrovski et al. 2017). Here we further investigated the separate contribution of each of these factors in more detail. Preparation of vials with processed food followed by larval transplantations was adapted from previous studies (Dombrovski et al. 2017, 2019, Fig. 1a). In our first experiment (Fig. 1b) we compared the effects of fresh versus processed food on larval pupariation and eclosion rates. To exclude the role of cooperative feeding, we used a low population density paradigm and transplanted 20 L2 wild type larvae (previously raised in normal conditions on fresh food) into plates and vials with processed food. We found that both pupariation (Fig. 1b) and eclosion (Fig. 1c) were significantly delayed in animals raised on processed food, but no difference was observed between rearing in plates and vials. Importantly, no effect on survival was found (Fig. S1a, left) and blind GMR-hid larvae displayed a similar eclosion delay in processed food. (Fig. S1b). In summary, processed food yielded in a ∼16h delay in pupariation and in eclosion in which a sub-cooperative number (20) of larvae were transplanted at L2 stage.

**Figure 1.**
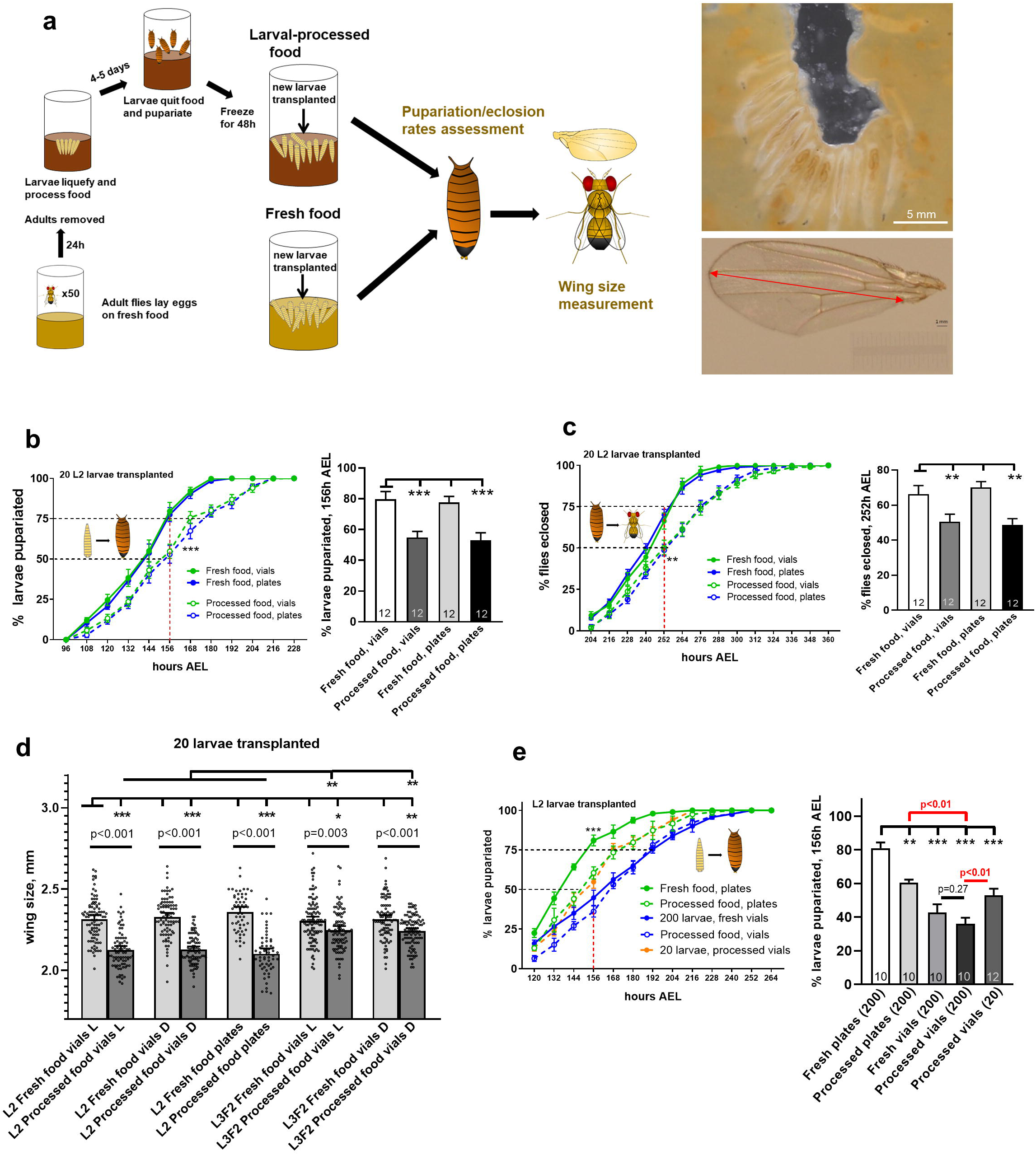
Processed food delays larval development and decreases animal size. **(A)** Schematic view of experimental procedures. In order to produce processed food, ∼50 adult flies were kept in fresh food vials for 24h and then removed, allowing a sufficient number of larvae to hatch and liquefy food within next 4-5 days. After all larvae pupariated, vials were frozen for 48h and cleaned. Newly collected larvae at a designated developmental stage were transplanted into defrosted vials with processed food in parallel with fresh food vials. Eclosion and pupariation rates were measured followed by assessment of adult female fly wing size to examine the effect of processed food on animal growth and development. **(B)** and **(C)** Processed food causes a significant developmental retardation. 20L2 wild type larvae were transplanted in vials or plates with fresh and processed food and subsequently evaluated for developmental rates. Rearing in processed food results in a consistent ∼16-20h delay in both pupariation **(B)** and eclosion **(C)**. No difference was seen between plates and vials. Percentage of larvae pupariated/eclosed was measured every 12h starting at 96 and 192h AEL, respectively. Here and further on, 156h AEL and 252h AEL checkpoints (indicated by dashed red lines) were compared for pupariation **(B)** and eclosion **(C)**, respectively, with data represented in bar graphs. **(D)** Processed food decreases wing size. 20 wild type larvae were transplanted into plates or vials with fresh or processed food with subsequent evaluation of adult fly wing size. In addition, effect of dark rearing or late transplantation (L3F2) was examined and compared between fresh and processed food vials. A significant ∼8-9% decrease in wing size was observed in L2 transplants raised in processed food, regardless of being reared in plates or vials and in normal light conditions (L) or darkness (D). A smaller 3-4% reduction in wing size was seen in L3F2 transplants raised in processed food, with no difference between normal light-dark regime and darkness. **(E)** Crowded conditions exacerbate developmental retardation. Pupariation rates were assessed in 20 and 200L2 wild type larvae transplanted into plates or vials with fresh or processed food. No difference was found between fresh and processed food vials. Importantly, 200 larvae raised in processed vials showed the biggest delay in pupariation, being significantly different from both 20 animals in processed food vials and 200 animals in processed food plates (highlighted in red). **See also Figure S1**

It has been reported that insufficient nutrition during larval stage reduces size in adult flies (Colombani et al 2003; Mirth and Riddiford 2007; Lavalle et al. 2008). Therefore, we next examined whether an observed developmental retardation was accompanied with size deficits. For this, we measured wing size in newly eclosed female flies, which serves a good estimate of general body size and weight (Taylor et al. 2008; Tang et al. 2011). As a control for experiments described in Figs. 2 and 3, we replicated the same conditions in the darkness. We found that, regardless of light regime and plates/vials animals reared on fresh food had significantly bigger wings (Fig. 1d), suggesting a negative impact of processed food on larval growth. In addition, we performed L3F2 larval stage transplantations into processed food vials and plates to see if decreased time spent in adverse nutritional environment would fully or partially rescue size deficits. We saw that, wing size in L3F2 transplants raised on processed food was significantly smaller compared to fresh food-reared larvae (Fig. 1d). However, the effect of processed food was less pronounced than in case of L2 transplants, indicating that time spent in processed food during larval stage negatively correlates with adult size. Alternatively, these results could also be explained by the fact that transplantation occurred after reaching critical weight (Mirth et al. 2005; Mirth and Riddiford. 2007, see Discussion for details). Survival rates were unchanged among all experimental paradigms (Fig. S1a).

**Figure 2.**
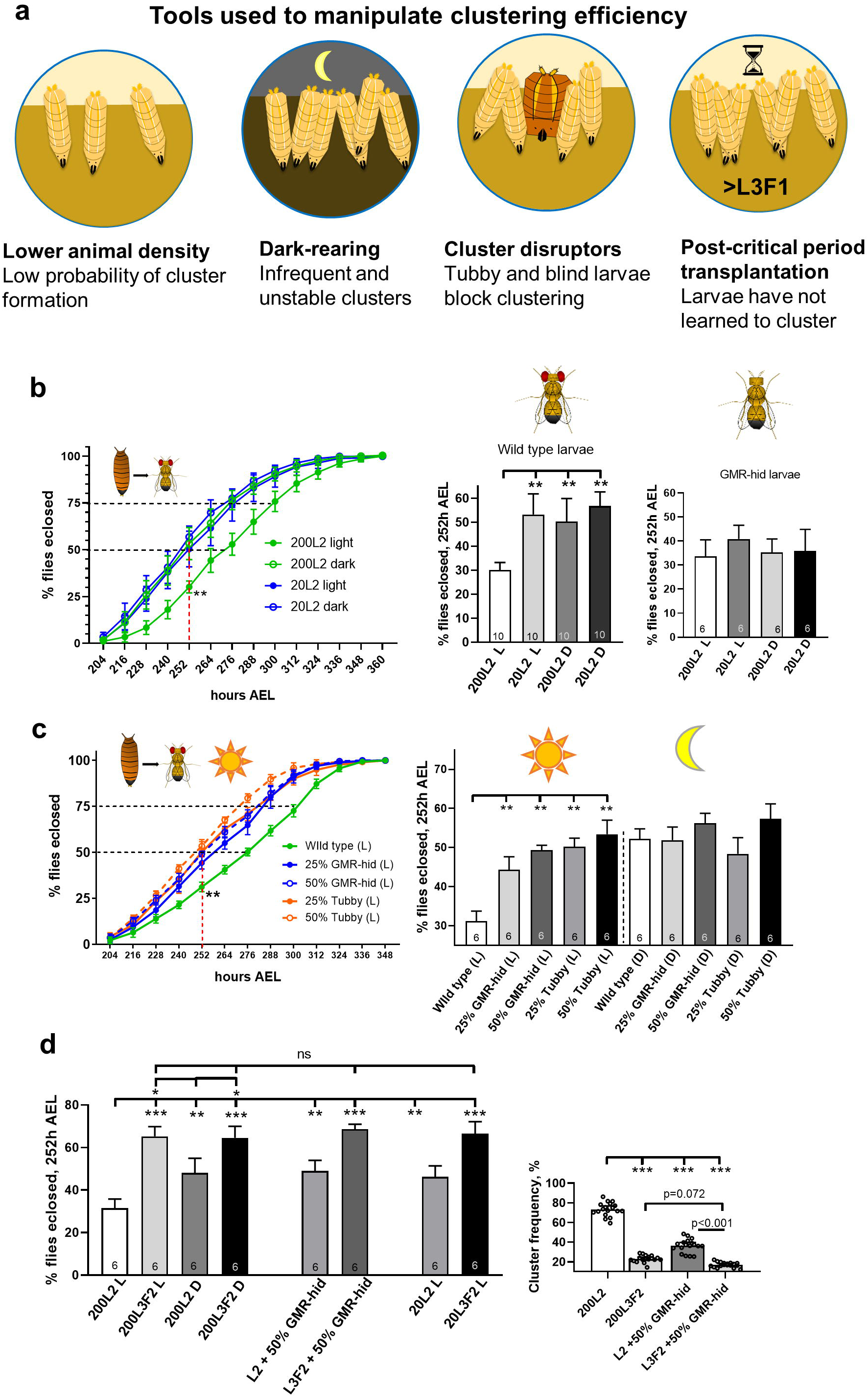
Cooperative foraging further delays larval development in processed food. **(A)** Schematic view of approaches further used (individually or in combination) to selectively reduce clustering efficiency in wild type animals. **(B)** A difference in developmental rates is seen between light- and dark reared larvae, but only in conditions that otherwise promote clustering. Eclosion rates were compared between 20 and 200L2 transplanted wild type larvae reared in normal light conditions (L) and in the darkness (D). Eclosion delay was significantly reduced in dark reared 200L2 larvae compared to animals raised in normal conditions. However, no significant difference was found in case of 20L2 transplants. In addition, no difference in developmental timing was seen among blind GMR-hid larvae exposed to the same experimental conditions (right panel). **(C)** Addition of cluster disruptors reduces larval developmental delay, but only in conditions that otherwise promote clustering. Eclosion rates were compared between 200L2 transplant groups containing 100% wild type larvae, 75% wild type + 25% GMR-hid or Tubby and 50% wild type + 50% GMR-hid or Tubby larvae. Same experiments were performed in the darkness. For animals reared in normal light conditions, cluster frequency was evaluated (right panel). For experiments performed in normal light conditions, addition of blind or tubby larvae resulted in a significantly decreased delay in eclosion compared to all-wild type groups. It was coupled with a corresponding decrease in clustering frequency. At the same time, no difference in eclosion rates was found between all-wild type and mixed groups for dark reared animals **(D)** Transplantation after the critical period for clustering reduces larval developmental delay. Eclosion rates were compared between wild type larvae transplanted at L2 and L3F2 stages, including comparison between 20 and 200 animals, normal light conditions (L) and dark rearing (D) and all-wild type populations and addition of 50% blind GMR-hid larvae. Regardless of any manipulations, L3F2 transplants displayed a significantly reduced delay in eclosion compared not only to 200L2 larvae in normal light conditions, but even 200L2s in the darkness and 20L2s as well as 200L2s in a mixed group (left panel). At the same time, L3F2 transplants showed significantly reduced clustering frequency compared to control wild type 200L2 and even mixed group 200L2 larvae (right panel). **See also Figures S1 and S2**

**Figure 3.**
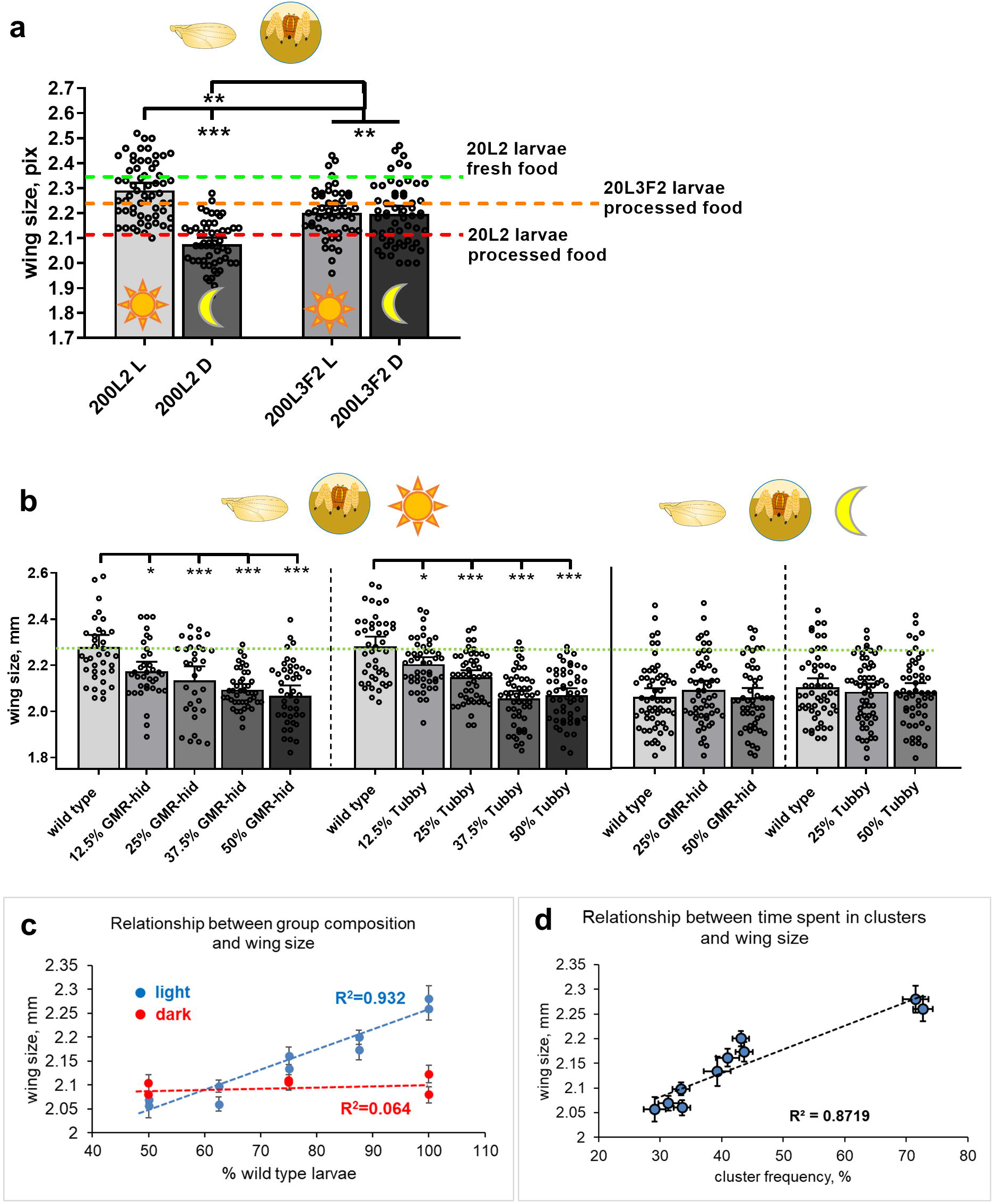
Clustering rescues fitness deficits caused by processed food. **(A)** Conditions that promote clustering also positively affect wing size. Wings size was compared between light- and dark reared L2 and L3F2 transplants (experiment shown in Figure 2D). 200L2 transplanted larvae reared in normal light conditions (L) have significantly bigger wings compared to their dark reared counterparts (D) and 20L2 larvae (indicated by a red dashed line, data from figure 1D) and only slightly smaller compared to 20L2 animals raised on fresh food serving as a positive control (indicated by a green dashed line, data from Figure 1D). In contrast, larvae transplanted at L3F2 stage display no difference in wing size between light- and dark reared animals, as well as 20L3F2 transplants in processed food (indicated by an orange dashed line, data from Figure 1D). Nevertheless, L3F2 larvae have smaller wings compared to 200L2 transplants reared in normal light conditions. **(B)** Group composition affects wing size, but only in normal light conditions. Wing size was compared between 200L2 transplant groups containing 100%, 87.5%, 75%, 62.5% and 50% wild type larvae reared in normal light conditions with the rest of the group comprising either GMR-hid or Tubby larvae in corresponding percentages (left panel). For dark reared animals (right panel), 100%, 75% and 50% wild type groups were used. A significant difference in wing size was found between all wild type animals from mixed groups and control 100% wild type groups raised in normal conditions. On the contrary, no difference in wing size was seen between groups of different composition in dark-reared larvae (wild type larvae from all groups had reduced wings compared to normal light reared all-wild type control group). **(C)** Relationship between group composition and wing size (data are related to Figure 3B). High positive correlation is seen between percentage of wild type larvae in a light reared group and wing size in flies derived from wild type larvae of the corresponding group. No correlation is observed in case of dark reared animals. Error bars represent SEM. **(D)** Relationship between time spent in clusters and wing size. Data are taken from experiments involving 200L2 wild type larvae and 12.5-50% mixed groups involving GMR-hid and Tubby larvae (results presented in Figure 3B, left panel). Clustering frequency is represented on the X-axis (data shown in Figure S2C). A significant positive correlation is seen between percentage of time a cluster was observed in a group of a designated composition and wing size in flies derived from wild type larvae of the corresponding group. Error bars represent SEM.

We next wondered how high population density changed the way processed food affects developmental timing especially with the appearance of cooperative foraging. For that, we used 200 L2 wild type larvae in vials and plates with either fresh or processed food and compared their pupariation and eclosion rates (Fig. 1e). For a control, GMR-hid larvae were exposed to similar experimental conditions (Fig. S1b). There was no difference in developmental rates between fresh and processed vials (Figs. 1e and S1c), which can be explained by very fast food processing by 200 animals. However, larvae reared in high-density in vials with processed food displayed dramatically delayed pupariation and eclosion compared not only to fresh food plates but also to processed food plates and processed food vials with low-density conditions (Figs. 1e and S1c). This is likely due to the formation of cooperative clusters and so this property was further examined.

### Cooperative foraging further delays larval development in processed food at high population density

In order to examine the role of cooperative foraging in larval development, we took advantage of approaches shown in Fig. 2a. In particular, we previously showed that wild type animals display dramatically reduced clustering when either deprived from light during specific periods of development (Dombrovski et al. 2019) or immediately after being placed in the darkness (Dombrovski et al. 2017, Fig. 2a). Here we compared developmental rates in wild type larvae reared in normal conditions and in the darkness and observed a significant delay in pupariation and eclosion times (Figs. 2b and S1d) normally reared animals compared to dark-reared counterparts. However, this effect was notable only in case of high population density that promotes clustering and no difference was found in case of 20 animals (Figs. 2b and S1d). Moreover, no difference in developmental timing was seen between normal- and dark reared GMR-hid larvae that cannot form clusters (Figs. 2b and S1b). Animal survival rates were not affected by dark rearing (Figs. S1b, S1d). Therefore, we isolated cooperative foraging as a single factor that contributes to developmental delay in otherwise unchanged conditions.

To further strengthen this notion, we used cluster “disruptors” as another tool to manipulate clustering frequency (Fig 2a), as our previous study indicates that introduction of non-clustering larvae into wild type foraging groups decreases cluster lifetime (Dombrovski et al. 2017). For this experiment, we compared developmental timing in all-wild type 200L2 larval groups and mixed groups (Fig. 2c) containing 25 and 50% GMR-hid or Tubby (Tb) larvae (both negatively interfering with clustering through their inability to either integrate into or efficiently cooperate within a cooperative group, as shown by Dombrovski et al. (2017). Vials were constantly recorded and cluster frequency in each vial was further assessed (Fig. 2c, right). The same experimental conditions were reproduced in dark reared larvae. We saw that the length of a delay in eclosion highly correlated with clustering frequency: it was most notable in all-wild type group and significantly decreased in a stepwise manner in 25% and 50% GMR-hid/Tb groups, similarly to a decrease in clustering frequency observed among the corresponding groups (Fig. 2c). Most importantly, no effect of group composition on developmental timing was seen in animals reared in the darkness (Fig. S2e, left), suggesting that clustering was a decisive factor. Survival rates were unchanged between light/dark conditions and group compositions (Fig. S2e, right).

Earlier studies also indicate that in order to cluster, larvae must pass through a visual critical period early in the third instar (Dombrovski et al. 2017, 2019) and animals transplanted into vials of processed food after this critical period show greatly reduced clustering. Therefore, we tested the effects of reducing clustering by post-visual critical period transplantation. We also reproduced the same experiment in the darkness, with 20 larvae and with a mixed group containing 50% GMR-hid larvae. We found that in standard high-density conditions, in light, L3F2-transplanted larvae displayed no visible delay in eclosion compared to L2 transplants (Figs. 2d and S2a). Interestingly, neither dark rearing, nor adding blind larvae and using 20 animals yielded a significant change in eclosion rates in L3F2 transplants. At the same time, these conditions had a strong impact on L2 transplants and shortened their developmental delay (Figs. 2d and S2a). Importantly, clustering frequency was significantly reduced in L3F2 transplants (Fig. 2d, right). These results implied that, since cooperative foraging was eventually absent in late-transplanted animals (Dombrovski et al., 2017), none of the factors reducing clustering efficiency was affecting their development, as opposed to L2-transplanted larvae being very sensitive to each factor (Fig. 2d). However, other interpretations of these results are possible. Increased survival rates compared to L2-transplanted larvae (Fig. S2b) and the fact that L3F2-transplanted blind GMR-hid larvae also showed a reduced developmental delay compared to L2 transplants (Fig. S2b) strongly suggested that the observed phenotype could also result from post-critical weight transplantation. In this case nutritional environment could have had no effect on L3F2 transplants’ developmental timing, thus making the effect of reduced clustering negligible. Therefore, this question required further clarification.

### Cooperative foraging during larval stage rescues size deficits caused by processed food in adult *Drosophila*

The experiments described above show that clustering further increases developmental time. We next examined how this delay affected adult animal size as measured by wing size. We first compared wing size in flies emerged from 200L2 transplanted wild type larvae raised on processed food and reared in normal light conditions and in the darkness (Fig. 3a), considering that light deprivation prevented animals from clustering. We found that wings of dark-reared animals were significantly smaller compared to the control group (Fig. 3a). Interestingly, a similar magnitude of difference in wing size was previously observed between 20 L2 transplanted wild type animals raised on fresh and processed food (Fig. 3a, dashed green and red lines, respectively). Thus, wing size was almost indistinguishable between animals raised on fresh food and animals derived from processed food, but only in conditions that promoted clustering (high population density and normal light regime). This suggests that clustering rescued the deficit in animal size caused by processed food. To further elucidate this phenomenon, we looked at how other factors interfering with clustering (Fig. 2a) affected wing size. Results from late post-critical period transplantation experiments of L3F2 larvae were intermediate between the two sizes in that wings were significantly smaller compared to positive control 200L2 transplants in light, they were also notably bigger than the negative control (200L2 in the darkness). However, they were almost identical to 20L3F2 transplants and not affected by light regime (Fig. 3a). This was in line with the data on L3F2 eclosion rates (Figs. 2d and S2a), implying that additional factors, such as post-critical weight effects play a role here (Mirth et al. 2005, Mirth and Riddiford, 2007, Mirth and Shingleton 2012).

In order to find more reliable correlation between changes in clustering and its effect on animal size, we compared wing size in wild type animals reared in clustering conditions but with cooperative behavior reduced in a regulated manner by the addition of cluster disruptors. (Figs. 2c and S1e). We added 12.5%-50% of either GMR-hid or Tb larvae to wild type animals transplanted at L2 stage. Clustering frequency in each group was assessed (as described in Dombrovski et al. 2017), and compared to the same conditions in darkness (Fig S2d). We found no differences in wild type wing size between groups of different composition reared in the darkness (Figs. 3b, right and 3c). At the same time, a clear relationship was seen between group composition and wild type wing size in light-reared animals (Figs. 3b, left and 3c): wing size decreased as clustering was suppressed (Fig. S2d). Overall, we were able to trace a strong positive correlation between the time wild type animals spent in clusters and their resulting adult size (Fig. 3d). In contrast, no relationship between clustering and wing size was seen in GMR-hid larvae (Fig. S2e), while the overall negative impact of processed food on wing size was present, consistent with the notion that blind animals are unable to use the benefits of cooperative foraging due to inability to form clusters.

## Discussion

Our study tests the idea that cooperative feeding among fruit fly larvae has a fitness benefit that is measured as body size and developmental time. Overall, we find mixed results in that the larval stage is increased in time but despite crowding, eclosed adults have normal size. Therefore, our study indicates that cooperative foraging in fruit fly larvae will be beneficial in some but not all conditions.

Processed food, which is predigested by larvae, slows developmental rates. It is likely that it has a lower nutritional value or at least an altered ratio of key macronutrients. According to a conventional notion, growth, larval developmental time and final size determination in insects including *Drosophila* are governed by of insulin-like hormones and TOR signaling in prothoracic gland (that is highly sensitive to nutritional status), as well as antagonistic actions and complex interplay of ecdysone and juvenile hormone (Colombani et al 2003; Layalle et al. 2008; Colombani et al 2012, Mirth and Shingleton 2012). In this context, malnutrition can have different impact on larval fate depending on whether it affects an animal before or after reaching critical weight, a key parameter that determines the readiness of the larvae to undergo metamorphosis and triggers the corresponding hormonal signals (Mirth and Shingleton, 2012). If occurring before that checkpoint in the middle of L3F1 stage, malnutrition only delays metamorphosis, but does not affect adult fly body size. Conversely, post-critical period starvation later in development has no influence on developmental rates, but dramatically reduces body weight and size (Mirth and Riddiford 2007; Mirth and Shingleton 2012). Interestingly, we observe both effects in solitary feeding animals, suggesting that processed food provides less nutrients, but not to an extent that would prevent larvae from reaching a minimum viable weight (Mirth and Riddiford 2007). In contrast, adult flies derived from clustering larvae lack size deficits, but display an even longer developmental delay. It implies that once animals engage in cooperative foraging groups, the efficiency of their feeding increases leading to a rescue in body size deficits. This idea is strengthened by our results showing that larvae transplanted into vials at L3F2 stage do not delay metamorphosis (explained by post-critical weight transplantation), but still have reduced body size, because they cannot cluster to feed more efficiently (transplantation occurs after critical period for clustering initiation, as shown by Dombrovski et al. (2017). Thus, cooperative foraging can be regarded as an evolutionary adaptation that outweighs malnutrition at the cost of developmental retardation. At the same time, specific mechanisms responsible for an additional delay in metamorphosis observed among clustering larvae remain unclear and are subject for future investigation.

A question arising from previous observations is how clustering enhances the efficiency of food consumption. Clustering larvae can take advantage of more efficient burrowing and reach fresher layers of food compared to solitary digging and non-clustering blind or socially naive animals (Dombrovski et al. 2017). This should also increase chances to avoid predation and infection by parasitoid wasps (Carton et al. 1985). In addition, clustering was shown to speed up the process of media liquefaction in vials (Dombrovski et al. 2017), that could in turn facilitate food ingestion by foraging animals. A more complex explanation features a phenomenon of communal exodigestion mostly observed among various fly larvae feeding on flesh and other high-protein substrates (Scanvion et al. 2018), but also documented in *Drosophila* (Gregg et al. 1990). Larvae are able to secrete a variety of enzymes digesting external polymers (amylose, cellulose and even chitin), therefore reducing energy expenditure per individual animal required to process and ingest a food source. Future metabolic studies are required to test this hypothesis.

In addition to the above, the influence of more complex and integrative factors on larval development in processed food and subsequent emergence of cooperative foraging is worth considering. As an example, it has been shown that gut microbiota in fruit flies is able to not only affect larval nutritional choices, feeding behavior (Venu et al. 2014; Leitão-Gonçalves et al. 2017; Qiao et al. 2019), and developmental rates (Shin et al. 2011), but also impacts kin recognition in adult flies (Lizé et al. 2013), which often is regarded as a source of social cooperation. Importantly, several studies suggest that kin recognition based on differences in gut microbiome community might as well be manifested during larval stage and thus determine animals’ ability to engage in cooperative foraging as larvae (Khodaei and Long 2019) and other complex social interactions during adult stage (Carazo et al. 2015), both of which are associated with significant fitness benefits. This may be especially relevant in our model system, where food processing by the larvae most likely leads to profound changes in the microbial composition. Therefore, future studies aimed to reveal connections between microbiome and cooperative foraging are required to shed more light on the evolution of social behaviors.

## Materials and Methods

### Fly stocks

Wild type Canton S (CS) flies were donated by Ed Lewis (Caltech), blind GMR-hid^G1^ strain was obtained from Bloomington Stock Center (#5771), w^-^;Sco/CyO and w-;sb/TM6B-Tb flies were kindly provided by Dr. Susan Doyle, University of Virginia.

### Fly stock maintenance and egg collection

All *Drosophila melanogaster* strains were raised in food vials (unless further specified in experimental details) containing standard Caltech food mixture (1000ml molasses, 14000ml H_2_O, 148g agar, 1000ml corn meal, 412g Baker’s Yeast, 225ml Tegosept, 80ml propionic acid) at 22°C, 30% humidity under standard 12/12h light-dark cycle (unless further specified in experiments involving dark-rearing) at ∼1000 lux light intensity. For egg production, ∼50 adult flies 3-4 days old were transferred into egg cups and kept in the same conditions. Eggs were collected on 35mm petri dishes containing standard agar-molasses food and yeast every 6 hours for experimental procedures and eggs/larvae were kept (unless further specified) in the same light/temperature/humidity conditions.

### Preparation of vials with “pre-processed” food and larval transplantations

Techniques of “pre-processed” vials production and transplantation of 200 L2 (60 hAEL) larvae were adapted from our previous studies (Dombrovski et al., 2017, 2019 and Fig. 1a). ∼50 adult flies 3-4 days old were kept in vials with fresh food for 24 hours and subsequently removed, vials were kept at standard conditions for 4-5 days allowing larvae to process food and pupariate. Vials were then frozen at -20°C for 48 hours, defrosted and cleaned from pupae before new larvae were transplanted into processed food. This approach was developed to minimize variance of “pre-processed’ food resources along with the absence of any unwanted animals and immediate exposure of transplanted larvae to designated food conditions

### Cluster frequency measurements

For cluster frequency measurements, vials with previously transferred 200 L2/L3F2 larvae of a designated genotype/condition raised in normal conditions were recorded for 5 consecutive days (24hrs non-stop) starting immediately after transplantation. Recorded videos were subsequently analyzed using ImageJ (32-bit version for Windows). Percentage of frames with a clearly observed larval cluster (defined as a group of 5 or more larvae aligned and oriented vertically and buried into the food for more than ¾ of the body length) was calculated for each 12-hour light period during 3 consecutive days of recordings for each individual vial (days 3-5 after transplantation for L2 larvae and days 1-3 after transplantation for L3F2 larvae) of a designated genotype/condition (each of these measurements represents a single data point on the corresponding graphs). Values were subsequently averaged and represented and mean values for each genotype/condition.

### Pupariation, eclosion and survival rates measurements

Newly formed pupae were counted on each food vial/plate of a designated genotype/condition twice a day (equal 12h periods and highlighted with a marker to avoid repeat counting. Eclosed flies were counted (for survival rates evaluation) and collected using CO2 anesthesia twice a day (equal 12h periods) and females were subsequently frozen at -20°C in 1.5mL plastic tubes for subsequent wing size measurements. For pupariation and eclosion measurements, percentage of animals reached a designated developmental stage was calculated relative to the total number of pupariated/eclosed animals counted by the final day of observations (not the total number of originally transplanted larvae). Survival values were estimated as ratios of eclosed flies to the total number of originally transplanted larvae. In case of mixed populations, only wild type CS flies were counted.

### Dark-rearing experiments

In all experiments involving light deprivation, dark-reared larvae were kept in a completely dark room for a designated time period in a food plate/vial additionally covered with a layer of aluminum foil. Dark rearing began immediately after larval transplantation at a designated stage and until the eclosion of all adult flies. Daily eclosion and pupariation rates measurements were performed in a room with dim red lights for each vial individually in order to minimize the time of possible light exposure.

### Wing size measurements

Wing size (which serves as an estimate of a body size) of previously collected and frozen females was measured using technique adapted from (Gilchrist and Partridge 1999) (distance from the base of the alula to the distal end of the third longitudinal vein, see Fig. 1a). A single wing from each animal was removed and mounted on a slide (with each slide representing 15-20 wings derived from animals from a single food plate/vial yielding in 4-6 slides per genotype/condition, where an individual wing measurement represented a single data point on the corresponding graphs). High-quality images of the slide were taken with a camera mounted on a tripod for subsequent wing size assessment using ImageJ (see below). Values were then averaged to give an estimate of the wing size for a designated genotype/condition.

### Photography and Video recordings

For cluster frequency analysis, videos were recorded on an iPhone 5 at full resolution and 1 frame/60” using “Lapseit” software for iOS. For wing images, a Nikon D3100 CMOS camera with 50mm lens and fitted with a Raynox Macroscopic 4x lens was used. Video analysis was further performed in iMovie and ImageJ (32-bit version for Windows).

### Statistical analysis

Unless otherwise stated, all data are presented as mean values and error bars represent 95% confidence intervals. Statistical significance was calculated by one-way ANOVA using Tukey’s method. When comparing two groups of normally distributed data, Student’s two-tailed unpaired T-test was used. *p<0.05; **p<0.01; ***p<0.001. Analysis was conducted using the GraphPad Prism 8 statistical software for Windows.

## Acknowledgements

We thank the Bloomington Stock Center for providing fly stocks. We thank Sarah Siegrist, Alan Bergland and members of the Condron lab for providing advice and comments. This work was supported Owen’s Family Foundation (BC) and the Jefferson Scholars Foundation (MD).

## Author contributions

Conceptualization & Methodology: BC, MD; Data collection: MD, RK, AM, HS; Data analysis: MD, RK; Writing, reviewing and editing: MD, BC; funding acquisition: BC, MD

## Supplemental Figure Legends

**Figure S1. Related to Figures 1 and 2**

**(A)** Survival rates are unaffected by food quality and rearing in plates/vials in case of 20L2 transplants (left panel, related to Figures 1B and 1C). No difference in survival rates was seen between normal and dark reared 20 L2 and 20 L3F2 transplants in fresh or processed food (right panel, related to Figure 1D).

**(B)** Processed food caused developmental retardation in GMR-hid larvae as well (left and middle panels, related to Figures 1B and 1C), but, unlike in case of wild type larvae, population size and light regime had no effect on developmental rates in processed food (left and middle panels, related to Figure 1E). Survival rates were not affected in any of those conditions (right panel).

**(C)** Eclosion rates in wild type larvae are also affected by crowded conditions (related to Figure 1E). Similar to pupariation (Figure 1E), an increase in animal number and transition from plates to vials significantly delay eclosion in processed food (left and middle panels). No significant difference was found between animals raised in fresh and processed food vials. Animal survival rates were not affected by any of the conditions mentioned above (right panel).

**(D)** Pupariation rates compared between 20 and 200L2 transplanted wild type larvae reared in normal light conditions (L) and in the darkness (D). Pupariation delay was significantly reduced in dark reared 200L2 larvae compared to animals raised in normal conditions, but no difference was found between light- and dark reared 20L2 transplants (left and middle panels, related to Figure 2B). Survival rates were not affected by population size and light regime (right panel).

**(E)** Addition of cluster disruptors does not affect eclosion rates if animals are reared in the darkness (left panel, related to Figure 2C). Eclosion rates were compared between 200L2 transplant groups containing 100% wild type larvae, 75% wild type + 25% GMR-hid or Tubby and 50% wild type + 50% GMR-hid or Tubby larvae. No difference in survival rates was observed among all-wild type and mixed groups in normal light conditions and in the darkness (right panel).

**Figure S2. Related to Figures 2 and 3**

**(A)** Post-critical period transplantation reduces eclosion delay in processed food (related to Figure 2D). Light regime does not affect eclosion rates in wild type L3F2 transplants compared to their L2 counterparts (left panel). Reduced animal density and addition of cluster disruptors has no effect on eclosion rates in L3F2 transplants compared to their L2 counterparts (right panel).

**(B)** Wild type L3F2 transplants show significantly higher survival rates compared to L2-transplnated larvae, regardless of animal density, light regime or addition of cluster disruptors (left panel, related to Figure 2D). GMR-hid L3F2 transplants similarly show increased survival rates compared to L2-transplanted blind larvae (right panel).

**(C)** Relationship between time spent in clusters and eclosion delay. A strong negative correlation is observed between cluster frequency and percentage of eclosed wild type animals at 252h AEL (data taken from experiments involving addition of cluster disruptors and L3F2 transplantations, related to Figures 2C and 2D).

**(D)** Cluster frequency is significantly reduced when cluster disruptors are added (related to Figure 3B). Sequential addition of 12%, 25%, 37.5% and 50% of GMR-hid or Tubby results in a gradual significant decrease in clustering frequency (top panel). A strong positive correlation is observed between percentage of wild type larvae in groups and clustering frequency (bottom panel).

**(E)** Controls for wing size in GMR-hid animals. Wing size is significantly reduced in animals reared in processed food compared to animals raised on fresh food. No effect of light regime was seen regardless of population size and transplantation stage. No effect of group size was observed. However, animals transplanted at L3F2 stage had slightly bigger wings compared to L2 transplants.

