## Supplementary figures and images for "Cooperative foraging behavior during larval stage affects fitness in adult *Drosophila*"

### Supplemental Figure 1

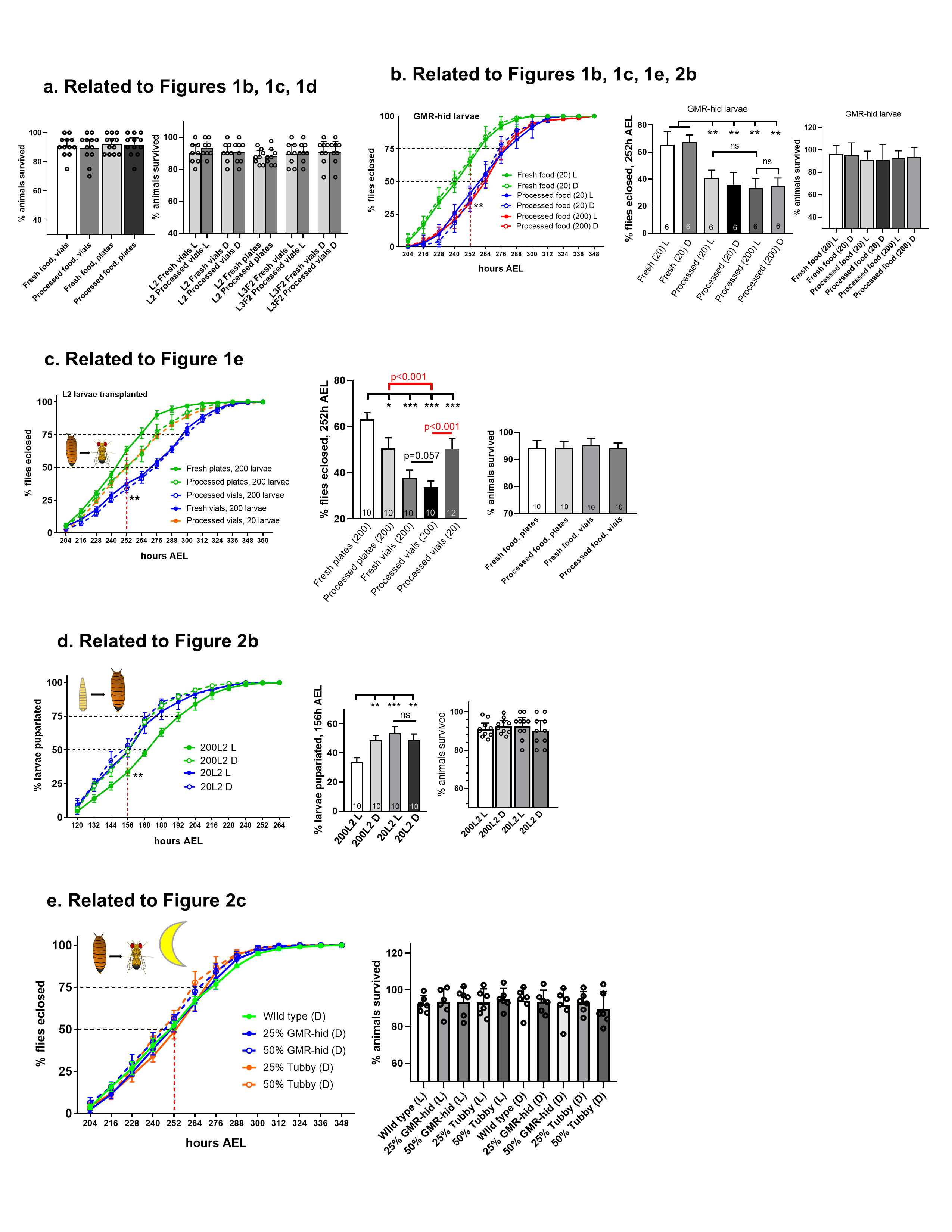

### Supplemental Figure 2

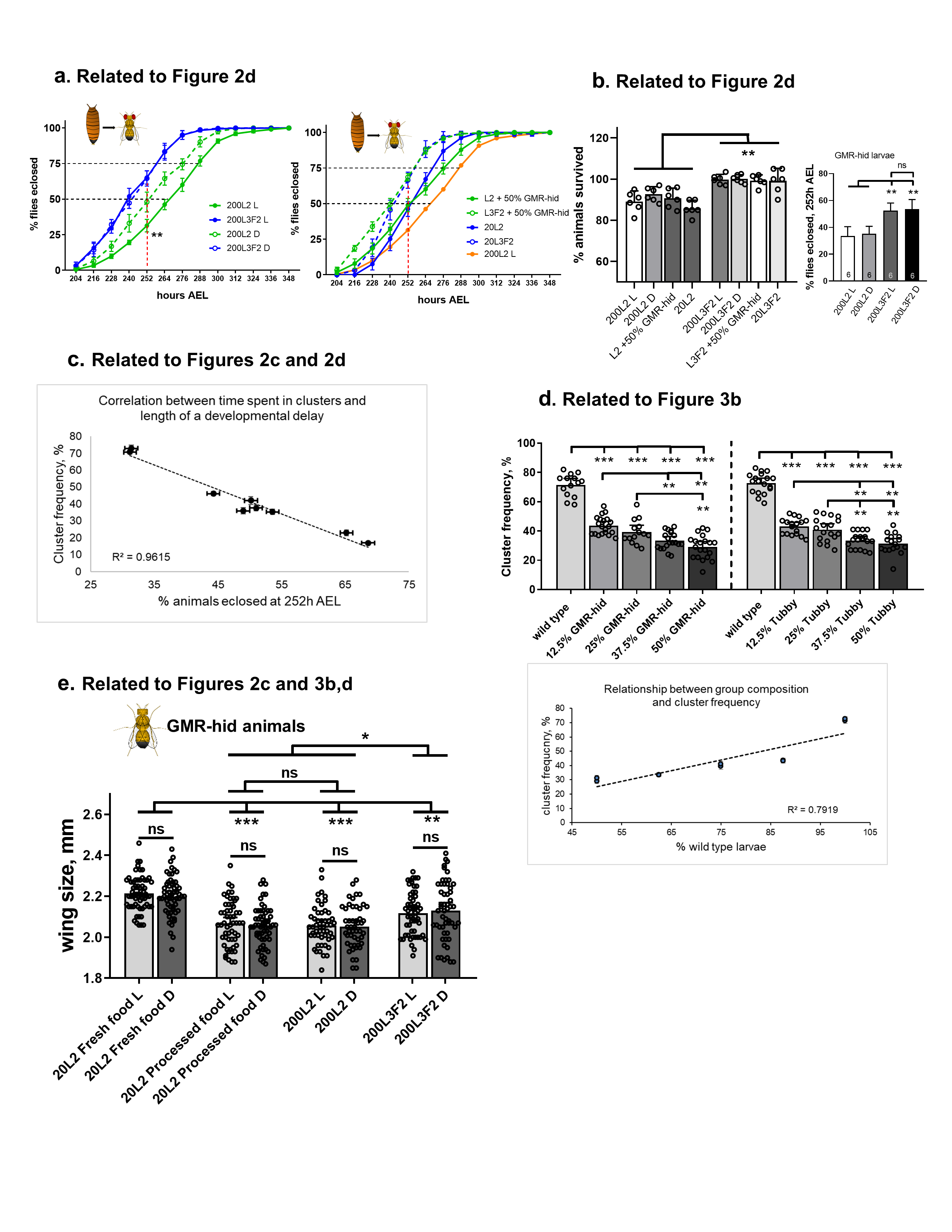
